# Measuring competitive exclusion in non-small cell lung cancer

**DOI:** 10.1101/2020.09.18.303966

**Authors:** Nathan Farrokhian, Jeff Maltas, Mina Dinh, Arda Durmaz, Patrick Ellsworth, Masahiro Hitomi, Erin McClure, Andriy Marusyk, Artem Kaznatcheev, Jacob G Scott

## Abstract

Therapeutic strategies for tumor control have traditionally assumed that maximizing reduction in tumor volume correlates with clinical efficacy. Unfortunately, this rapid decrease in tumor burden is almost invariably followed by the emergence of therapeutic resistance. Evolutionary based treatment strategies attempt to delay resistance via judicious treatments that maintain a significant treatable subpopulation. While these strategies have shown promise in recent clinical trials, they often rely on biological conjecture and intuition to derive parameters. In this study we experimentally measure the frequency-dependent interactions between a gefitinib resistant non-small cell lung cancer (NSCLC) population and its sensitive ancestor via the evolutionary game assay. We show that cost of resistance is insufficient to accurately predict competitive exclusion and that frequency-dependent growth rate measurements are required. In addition, we show that frequency-dependent growth rate changes may ultimately result in a safe harbor for resistant populations to safely accumulate, even those with significant cost of resistance. Using frequency-dependent growth rate data we then show that gefitinib treatment results in competitive exclusion of the ancestor, while absence of treatment results in a likely, but not guaranteed exclusion of the resistant strain. Finally, using our empirically derived growth rates to constrain simulations, we demonstrate that incorporating ecological growth effects can dramatically change the predicted time to sensitive strain extinction. In addition, we show that higher drug concentrations may not lead to the optimal reduction in tumor burden. Taken together, these results highlight the potential importance of frequency-dependent growth rate data for understanding competing populations, both in the laboratory and the clinic.

## Introduction

Given our current understanding of intratumoral heterogeneity, treatment resistance after continuous dose chemotherapy is an expected consequence. Genomic instability^1^, inherent to the development of most cancer^2–5^, results in the accumulation of a variety of aberrations within a single tumor population.^6^ While only a small subset of these randomly distributed changes will contribute directly to driving carcinogenesis, this diverse population comprised of phenotypically distinct subclones results in increased resilience of the overall tumor population across a wide range of external stressors.^7–9^

These distinct subclones do not live, grow, or reproduce in isolation. With this diverse cellular population comes a diverse range of intercellular interactions^10,11^. Complex systems cannot often be fully described empirically, and their dynamics can be difficult to intuit from measurements of their parts. In these situations, mathematical models have historically played a role. Specifically, evolutionary game theory has been effective in predicting the evolutionary consequences of interactions in large multicellular ecosystems, such as fisheries^12^ and game reserves.^13^ More recently, these evolutionary game theoretical models have been utilized to gain insight into phenotypic shifts that occur within tumor ecosystems.^11,14–17^ As a consequence, we must understand both the absolute fitness advantages of particular subpopulations in the selecting environment (monoculture), as well as how competing clones modulate that advantage as a function of population frequency (co-culture).^18^ This frequency-dependent growth can transform the power of evolution^19^ and acts to shape treatment-naïve tumor ecosystems and influences inevitable development of resistance in post-treatment environments.^20–23^ As traditional treatment protocols continue to fail, more evolutionary-based treatments that rely on judicious treatment schedules and cooperative dynamics between populations have gained in popularity.^10,11,24–31^

Dynamic therapeutic protocols using models of this type have already made their way into the clinic with promising results.^27^ While this highlights the value of game theoretical models for treatment optimization, the specific model in this, and other clinical trials have been selected and parameterized mainly based on biological conjecture and intuition.^32,33^ Instead, for each clinical condition, a different model and parameters would likely be needed to accurately capture intratumoral dynamics. As such, reproducibility of this initial success across different tissues and environmental contexts is contingent on our ability to measure subclonal interactions in the lab prior to transitioning to clinical practice. These interactions can greatly influence the evolutionary trajectory of the tumor; therefore, incorrect characterization could unintentionally worsen treatment outcomes.

One such concept that is frequently assumed to be the driver of inevitable treatment resistance within these models is that of competitive release.^34^ This phenomenon was first described by Joseph Connell while studying the distribution of barnacles off the shore of Millport, Scotland, where it was observed that two species occupied two distinct horizontal zones on the shoreline.^35^ Connell determined that the upper species, *Chthamalus stellatus*, was competitively excluded from populating the lower region as a result of competitive interactions with the lower species, *Balanus balanoides*. Experimental removal of *Balanus* by Connell released *Chthamalus* from this competitive exclusion, which resulted in expansion into the lower horizontal zone. Similarly, in tumors, it is thought that selective killing of sensitive cells during therapy removes competitive restrictions on resistant populations, allowing for their outgrowth and subsequent therapeutic failure. While intuitive in theory and observed in bacteria^36^ and parasites^37–39^, empiric evidence of the dynamics that underlie this phenomenon in cancer have, to our knowledge, yet to be elucidated.

As a population becomes increasingly resistant to a new treatment, it is common for that population to pay a ‘fitness cost’ to maintain that resistant mechanism, leading to a reduced growth rate when compared to the ancestor from which it was derived. This has led many researchers to suggest that the sensitive ancestor is likely to out-compete the resistant clone when selection is removed, and thus treatment holidays may be beneficial to the maintenance of a treatable cancer population^27,40^. In some cases the fitness cost may be significant enough to result in competitive exclusion of the resistant strain upon treatment withdrawal. Instead, in this work we consider the importance of empirical, frequency-dependent growth rate measurements. Beginning with PC9, a model cancer system for EGFR TKI resistance in NSCLC, we show that competitive exclusion requires one population to out-compete (have a higher growth rate than) another population under all possible population frequencies. We then measure these frequency-dependent growth rates for a gefitinib-resistant population and the ancestor from which it was derived and show that competitive exclusion is likely, but not guaranteed. We then show that the addition of gefitinib shifts the frequency-dependent growth rates such that the resistant strain will competitively exclude the ancestor at all tested concentrations. Finally, combining our empirically derived growth measurements with traditional competition simulations, we demonstrate that the inclusion of ecological effects can significantly alter the predicted time to exclusion of the ancestor, and thus alter the time required to reach an untreatable resistant tumor. In addition, we show that contrary to the maximal tolerable dose hypothesis, higher drug doses may not constrain tumor burden better than lower doses for our system.

### Box 1

Evolutionary Game Assay

#### Tracking individual subclones in heterotypic cultures

To track differential growth dynamics of two populations in the same culture, each population was transduced with a vector encoding a different heritable fluorescent protein. For this experiment, the resistant and parental cells were made to stably express mCherry and EGFP respectively. The expression of these proteins was linked to nuclear localization signal (NLS) repeats for localization of the fluorescent signal into each cell’s nuclei. This increases resolution and accuracy of cell number counts at higher confluency. Once plated together in heterotypic culture, each subclone could be tracked through time in their respective fluorescent channel using time-lapse microscopy systems (**Fig 1A**).

**Figure 1.**
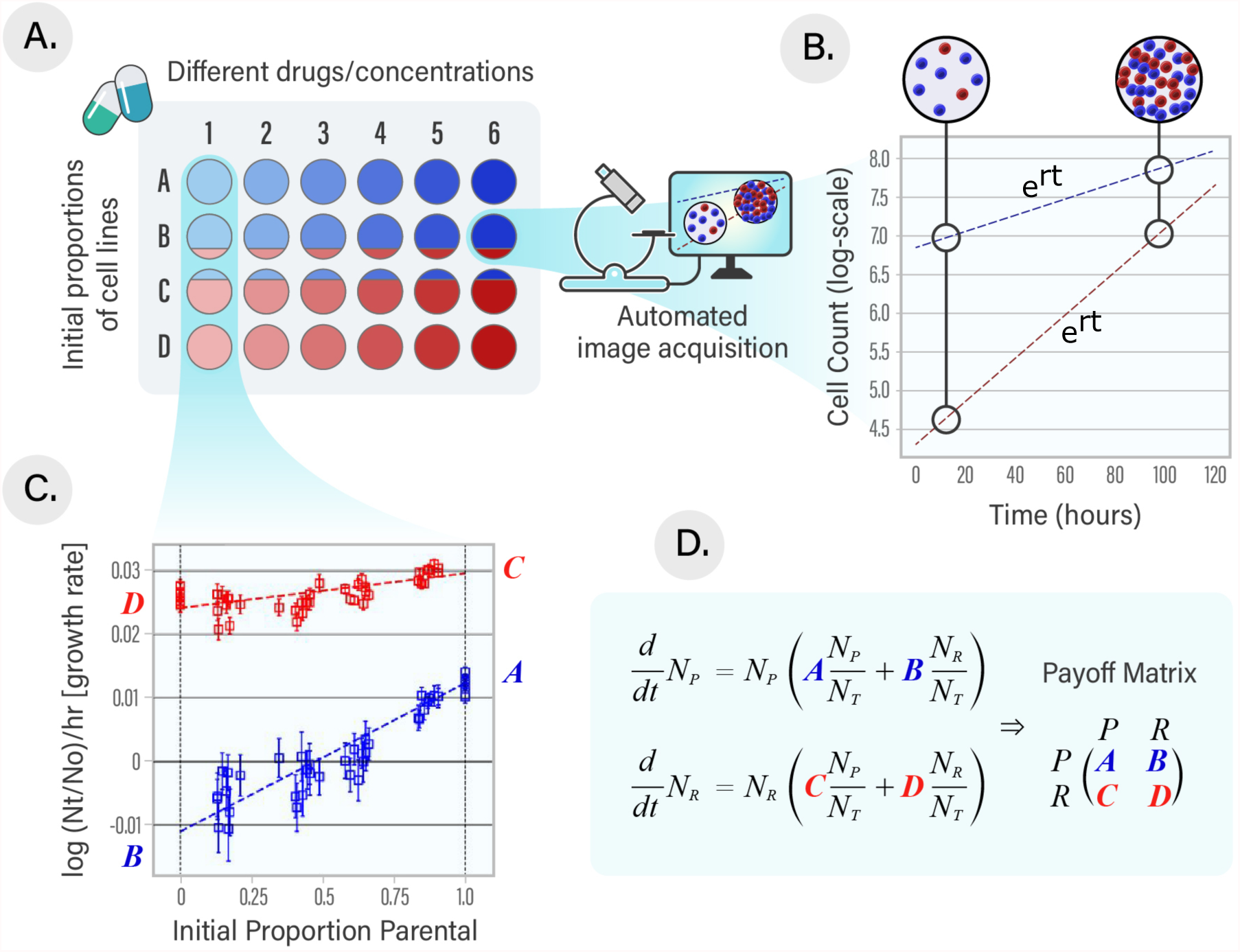
Experimental design of measured evolutionary games. **(A)** A Gefitinib-resistant cell line (red) and its sensitive ancestor (blue) were plated across a full spectrum of initial proportions (A, B, C, D…) and in a range of different different drugs and concentrations (1, 2, 3, 4…) in a 96 well plate. **(B)** Automated time-lapse microscopy imaging captures the composition of the population in each well every four hours. The cell lines were labeled via lentiviral transduction in order to precisely quantify the growth rate of the two separate populations. Cell number counts were extracted from each fluorescent image and plotted against elapsed time to derive growth rates in each well for each population. **(C)** Extracted growth rates were then plotted as the frequency-dependent growth rate of each population (growth rate as a function of ancestor population fraction). **(D)** Evolutionary game dynamics are quantified (fitness functions and associated payoff matrices) using least squared regression and intercepts of p= 0 and p= 1.

#### Translating image information into growth rates

Cell number counts were extracted from each fluorescent image at each time point throughout the time series. Exponential growth rates where determined via semi-log regression of change in cell number against change in time (hours) using the Theil-sen estimator (**Fig 1B**).

#### Fitness functions - growth as a function of population composition

To find the dependence of fitness on the frequency of subclonal interaction, least squares regressions were performed on the growth rate against the initial proportion of parental in each well (**Fig 1C**). This regression was weighted against the inverse of the errors 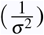 associated with each growth rate. The resulting linear equations describe growth as a function of the initial proportion of the opposing subclone:

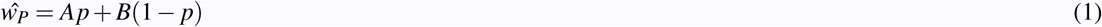

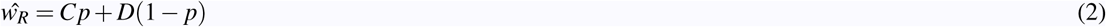

#### Game theoretical payoff matrix

To clearly represent the fitness outcome of specific interactions, payoff matrices corresponding to each of the different conditions can be derived from the resulting fitness functions (**Fig 1D**) For example, the fitness outcome of parental cells interacting with one another occurs when *p* = 1, which translates to *ŵ*_*P*_ = *A*. Similarly, the fitness outcome of when parental interacts with resistant occurs when *p* = 0, which translates to *ŵ*_*P*_ = *B*.

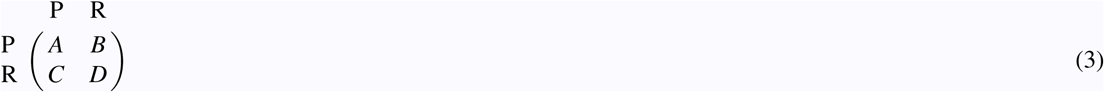

## Results

### Ecological interactions ameliorate some, but not the entire fitness cost associated with resistance, resulting in competitive exclusion of the resistant population in the absence of treatment

We investigated the evolutionary games (frequency-dependent ecological interactions) between ancestral and gefitinib-resistant cell lines in the lung adenocarcinoma cell line PC9. To create the resistant line, we exposed the population to 1*µ*M gefitinib for 6 months. In parallel, the original ancestral PC9 cell line was propagated in a matched volume of DSMO for 6 months. A high initial dose of gefitinib was chosen to select for pre-existing resistant populations rather than drug tolerant subclones.^41^ We then used the game assay we have previously developed to quanitfy ecological interactions between populations.^30^. Briefly, we co-cultured the derived gefitinib-resistant cell line, with the DMSO-propagated ancestral cell line at varying ancestral population to resistant population ratios (**Fig 1A**) in a 96-well plate. Using an automated incubator (BioTek Biospa) and time-lapse microscopy setup, we imaged the wells every 4 hours. Cell lines were transduced with EGFP and mCherry fluorescent proteins that, when combined with image processing software, allowed for quantification of population-level cell counts, and therefore population growth rates (**Fig 1B**). Then, by combining the parallel experiments done at varying initial parental (ancestor) populations, we can plot the frequency-dependent growth rates for both the resistant and parental cell lines (**Fig 1C**). Finally, we can extract the payoff matrix that describes the evolutionary game dynamics as shown in **Fig 1D**.

In addition to the frequency-dependent growth rate measurements based on heterogeneous (mixed parental and resistant populations), we also performed the more standard measure, of the homogeneous, monotypic growth rates for both cell lines in DMSO. In this case, there is a substantial growth cost to the resistant phenotype, which grows at roughly three-fourths (75.6%) the rate of the monotypic ancestral PC9 population (**Fig 2B**, left panel). Fitness costs are often assumed in treatment resistant populations of EGFR driven NSCLC,^42^ however this feature may not be generalizable across all NSCLC types.^30^ While it is tempting to extrapolate this data and suggest these growth rate differences necessitate competitive exclusion of the resistant cell line due to its lower growth in DMSO, this is not necessarily so. Ecological interactions between the populations can ameliorate the fitness cost associated with resistance as shown in the left panel of cartoon plot, **Fig 2A**. The difference between the resistant and ancestor population’s frequency-dependent growth rate is known as the gain function. For competitive exclusion to occur, the value of the gain function cannot change sign for any population frequency. That is, if the gain function is positive (or negative), it must remain positive (or negative) for all population frequencies. If instead the gain function transitions from negative to positive, or vice versa, a fixed point will occur.^43^ If the fixed point is stable (negative to positive gain) then it will allow for co-existence of both populations, otherwise it will be a bifurcation for the population’s dynamics. Our experiments reveal the ancestor out-competes the resistant population at all population frequencies, likely resulting in a complete competitive exclusion of the resistant population (**Fig 2B**, right panel).

**Figure 2.**
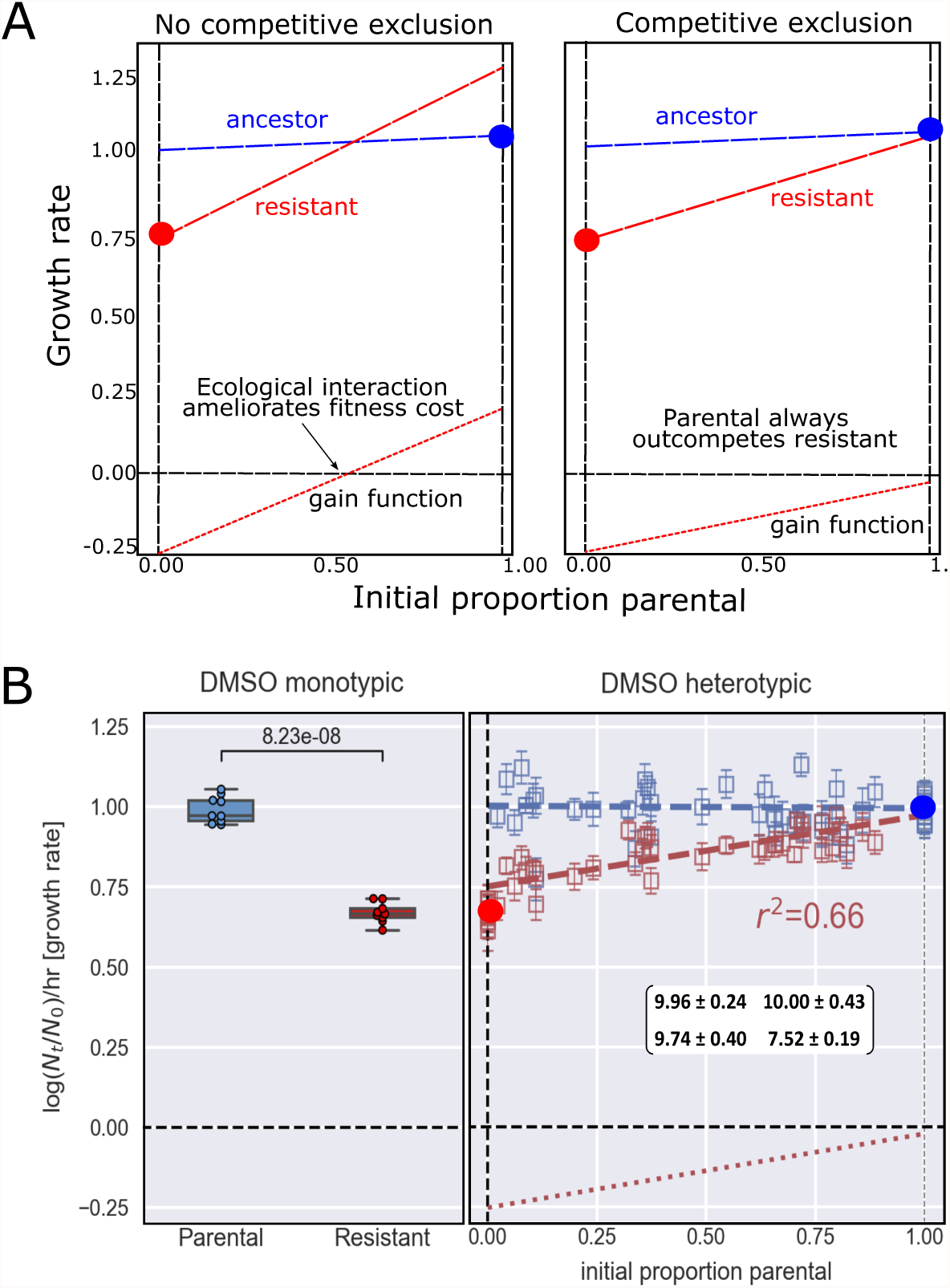
Ecological interactions alter but do not ameliorate the resistant clones fitness cost and result in competitive exclusion of the resistant strain in DMSO. **A**. Monotypic growth measurements are insufficient to predict competitive exclusion. The left and right panel both depict a resistant strain with a significant fitness cost associated with its resistance. Red circle at *p* = 0 represents monotypic resistant growth, while the blue circle at *p* = 1 represents monotypic ancestral growth. **Left Panel:** We see the ecological interaction is large enough to overcome the fitness cost when co-cultured with a majority parental population, resulting in no competitive exclusion. **Right Panel:** While there is significant ecological interaction, it is insufficient to overcome the fitness cost, resulting in complete competitive exclusion. **B. Left panel:** Monocultures in DMSO shows significant difference in growth between subclones (*p <<* 0.001), highlighting the cost associated with the resistant phenotype. **Right panel:** Heterotypic cultures in DMSO reveal strong frequency-dependent interactions that modulate the resistant populations growth, however these ecological forces are insufficient to completely overcome the monoculture fitness cost, leading to competitive exclusion of the resistant subclone. The ancestor’s growth remains consistent at all frequencies. Plotted values were normalized against mean monotypic parental growth in DMSO. Values in displayed game matrix have been scaled by a factor 10 for ease of comparison.

Interestingly, however, the fitness cost of the resistant population is almost entirely ameliorated by ecological interactions occurring at high ancestral population frequencies. Because this interaction occurs at resistant population fractions near zero that are hard to reliably measure empirically, it is possible that the resistant population is not completely competitively excluded, and a fixed point may exist at this extreme. If this were to be the case, it could highlight one potential way a sensitive population may generate and maintain drug-resistant populations without losing population-level fitness, as the resistant strain with a significantly fitness cost could be maintained at low population frequency in the absence of drug.

### The addition of drug switches which population is competitively excluded

Next we sought to quantify the ecological interaction between the parental and resistant cell lines under the application of increasing gefitinib concentrations (**Fig 3A**). Interestingly, we see that as the concentration of gefitinib increases, the slopes of the frequency-dependent growth rates (and therefore, ecological interaction magnitude) also increases. We can quantify this more clearly by instead plotting the gain function, or the difference between the growth of the ancestor and resistant cell lines (**Fig 3B**). When visualized this way, it is immediately apparent that under treatment of DMSO the resistant strain is competitively excluded by the ancestral strain (blue line is completely above x-axis). In addition, we can conclude under the treatment of all tested gefitinib concentrations the cell line that is out-competed is reversed, and the ancestral strain is competitively excluded (all other lines are completely below the x-axis). Finally, we can see the slope changes from negative for DMSO to increasingly positive as the concentration of gefitinib increases. This depicts an increasing ecological interaction strength.

**Figure 3.**
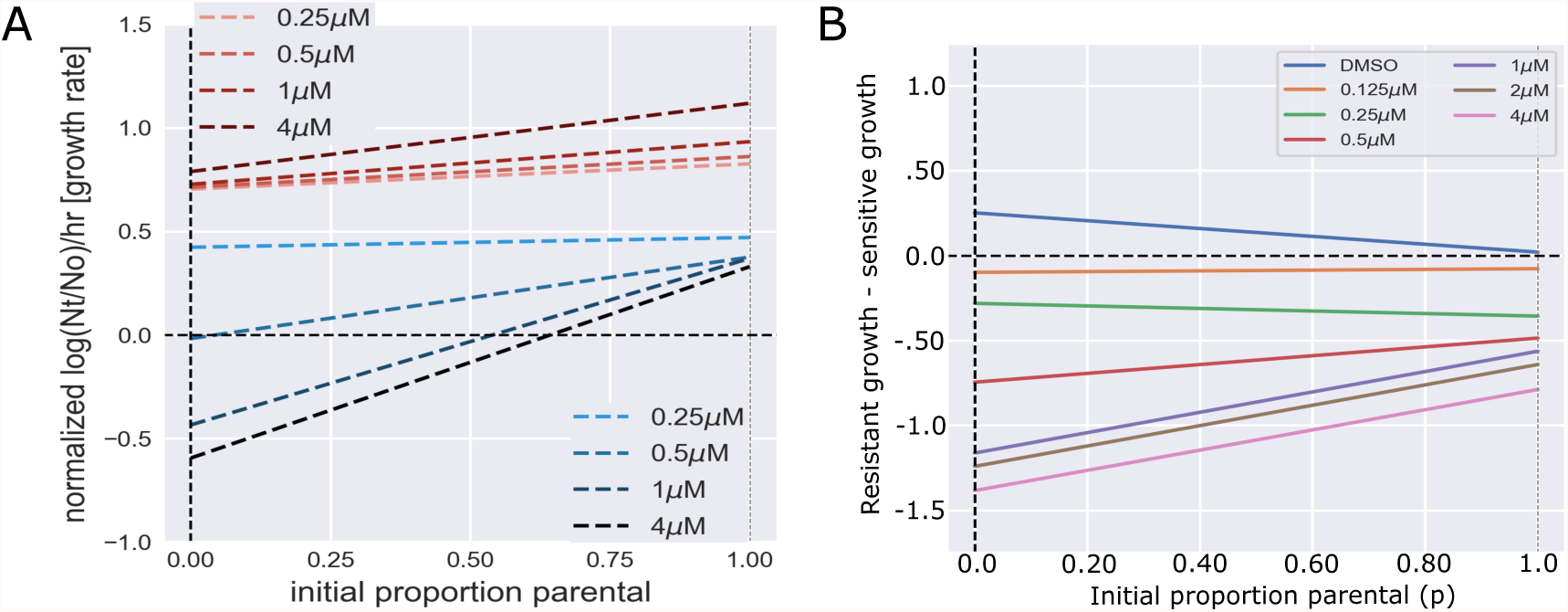
Increasing gefitinib concentration switches which population is competitively excluded from resistant to ancestor. **A**. Frequency-dependent growth rates were measured for several concentrations of gefitinib for both gefitinib-resistant (red) and the sensitive ancestor (blue). Growth values were normalized against the ancestor’s mean monotypic growth in DMSO. Under all concentrations tested we see the resistant strain out-competing the ancestor at all population fractions, resulting in competitive exclusion of the ancestor. **B**. Gain function (growth of resistant cell line - growth of ancestral cell line) for all seven tested concentrations of gefitinib. DMSO’s (blue) gain function exists entirely above *y* = 0 signifying competitive exclusion of the resistant cell line. For all non-zero concentrations the game dynamics shift to exist entirely below *y* = 0, indicating competitive exclusion of the ancestral strain.

### Competition simulations reveal importance of ecological effects in ancestral extinction rates and tumor burden calculations

We built two kinds of mathematical models to explore the outcomes of ancestral extinction rates and tumour burden further. We extrapolated the derived fitness functions out through time using both replicator dynamics (**eq. 9**) and a practical derivative of the Lotka-Volterra (LV) equation (**eq. 11**) that allows for competitive exclusion of interacting species and better modelling of the timescales of extinction dynamics^44^. Both of these models attempt to predict population trends through time. The replicator dynamics does this while modelling only frequency change and not population size^32^. The LV model constrains the population to a strict user-chosen carrying capacity or maximum size. *In vivo* tumor growth likely falls somewhere in between, as nutrient availability and space constrains growth in a more fluid manner through mechanisms such as angiogenesis. For each model, relative time to extinction was determined for the range of doses, where extinction is defined as proportion of the population, *p*, falling below *<* 0.01. As expected, both models predict faster extinction of the sensitive population at higher doses (**Fig 4A,B**).

**Figure 4.**
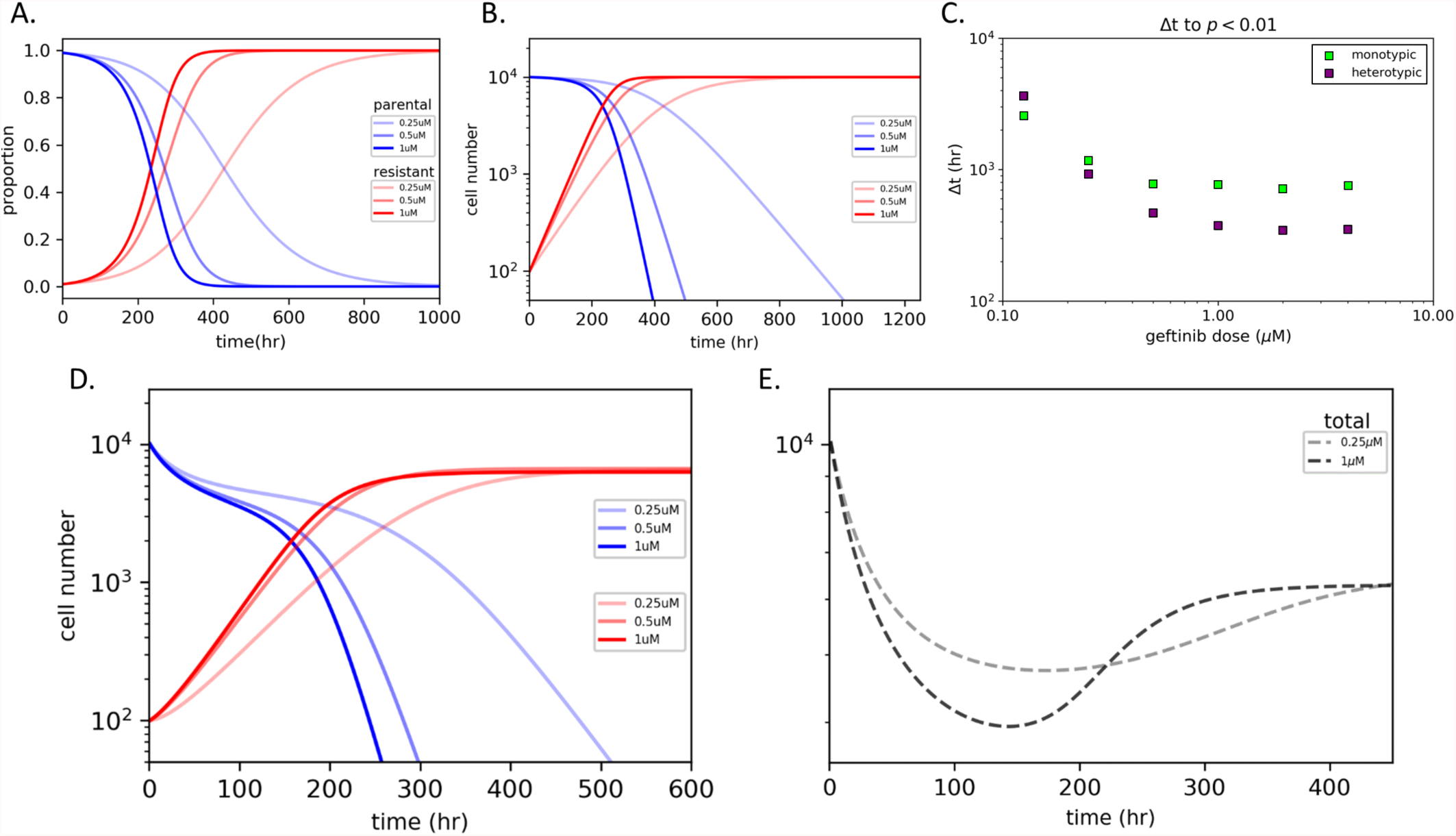
Evaluation of growth models with empirically derived parameters highlights rapid acceleration of competitive release via non-cell autonomous interactions and demonstrates persistence of qualitative features across the spectrum of models tested. The same initial parameters were used for each model (*p* = 0.99). **A**. Replicator dynamics showing proportional shifts of both competing cell populations over time in three gefitinib doses. **B**. Lotka-Volterra (LV) model of outgrowth in constrained environments with equal carrying capacities (*K*_*p*_ = *K*_*r*_). **C**. Time to extinction (defined as the proportion of the population, *p*, dropping below 0.01) across a range of gefitinib doses was determined and compared between cell autonomous (monotypic) and non-autonomous (heteorotypic) growth. **D**. LV model with unequal carrying capacities (*K*_*i*_ = *K*_*max*_*α*_*i*_ where *α*_*i*_ = *r*_*mono*_*/r*_*max*_). **E**. Estimates for changes in total tumor burden for relative LV model in 0.25*µ*M and 1*µ*M of gefitinib. Treatment with the lower dose of 0.25*µ*M had a smaller initial response to therapy, but longer overall response due to delayed parental extinction and maintenance of heterogeneity over a longer period of time.

While LV models can be quite sensitive to user-chosen carrying capacities, when the carrying capacities of both subclones are equal (*K*_*p*_ = *K*_*r*_ = *K*_*max*_), heterogeneity was maintained at identical time scales when compared to the replicator equations. To evaluate the impact of frequency-dependent growth, the results were contrasted between models run with monotypic culture growth parameters and those measured in heterotypic cultures (**Fig 4C**). Interestingly, the model predicts that as drug concentration increases, the monotypic growth rates increasingly overestimates the time to extinction of the ancestor population when compared to the more accurate heterotypic growth data. This is because the heterotypic data captures the accelerated rate of competitive exclusion that occurs as a result of ecological effects. That is, the difference between the resistant and parental growth rates is larger at every population fraction than the measured monotypic growth rates.

Because the assumption of equal carrying capacity may not be true of *in vivo* contexts, we varied the relative carrying capacity of the two population to be a ratio of the monotypic growth rates, scaled by their maximum rate in the absence of treatment (**Fig 4D**). Promisingly, these results are qualitatively identical to the results gathered from using equal carrying capacity LV models, and those predicted from replicator dynamics. However, clinical tumor burden is likely most correlated with the total tumor size, and not the size of a particular sub-population. With this is mind we show sample time traces of total tumor population over time in response to two drug concentrations (**Fig 4E**). Interestingly, we observe that while a lower drug concentration may lead to a smaller initial tumor decline than a larger drug concentration, the lower concentration leads to a prolonged heterogeneous, and therefore more sensitive, tumor state. This is the result of a lower dose of gefitinib prolonging the competition between ancestor and resistant populations, not allowing for a competition-free expansion by the resistant sub-population.

## Discussion

Our work provides an extensive quantitative study of the frequency-dependent interactions between an experimentally derived gefitinib-resistant PC9 cell-line and its sensitive ancestor. We have shown that a fitness cost resulting from resistance may be insufficient to result in competitive exclusion of the resistant population in the absence of drug. Instead, frequency-dependent ecological interactions with the ancestral population may ameliorate the fitness cost, leading to a potential safe harbor for small resistant populations. As a result, future studies focused on competitive exclusion would benefit from an examination of frequency-dependent ecological interactions.

In addition, our work also examined how the ecological interactions may shift under increasing gefitinib doses. Our results show a shift from competitive exclusion of the resistant population to a competitive exclusion of the ancestral population as gefitinib dose is increased. Then, with simulations we demonstrate that the inclusion of ecological effects can significantly alter calculations of ancestral extinction rate and temporal tumor burdens. Past work has shown how drug and tumor microenvironments can fluctuate in space^45–47^, suggesting that these ecological interactions are likely to fluctuate over space as well. Future experimental and modeling work will attempt to untangle these potential spatial contributions.

In interpreting our work several limitations are to be kept in mind. Measuring frequency-dependent ecological interactions is extremely challenging. As such, our intention was not to produce a highly accurate, clinical model of ecological interactions that can inform current cancer treatments. Instead, we focus on measuring the ecological interaction between two competing populations, and how that may influence *in vitro* laboratory experiments. In beginning to understand this more idealized scenario we hope to understand more general principles that can be tested in more complex models of resistance and ecological interactions. With that said, it is our hope to build on this simple model to account for more complex interactions.

While our work compliments recent studies on competitive exclusion and evolutionary game theory, it also raises interesting new questions for future work. For example, clinical tumors are highly heterogeneous. Extending this work to include three^48^ or more types will allow us to better model more clinically relevant resistance evolution. Finally, our results suggest that ecological effects are an important consideration in competition experiments, and continuing to show this empirically remains a priority going forward.

## Methods

### Cell lines

All cells were cultured in Roswell Park Memorial Institute (RPMI) media supplemented with 10% fetal bovine serum (FBS) and 1% penicillin/streptomycin.

Parental and resistant cell lines were established from the same initial population of PC9 cells (Sigma-Aldrich 90071810). Resistant population was cultured in 1uM of gefitinib (Cayman 13166) for greater than 6 months, until a population of stably growing cells was observed. Resulting subpopulations exhibited noticeable visual morphological differences in culture. The parental population was cultured in parallel in matched volumes of dimethyl sulfoxide (DMSO) (Sigma-Aldrich 276855) for the same duration as a vehicle control.

Resulting resistant and parental subclones underwent lentiviral transduction with plasmid vectors encoding EGFP- and mCherry-fluorescent proteins with attached nuclear localization sequence (plasmids were a gift from Andriy Marusyk’s lab at Moffitt Cancer Center). Derivative cell lines with heritable fluorescent protein expression were selected for in puromycin (MP Biomedical 100552).

### Experimental design

Cells were harvested at 70-80% confluence, stained with trypan blue (Corning 25-900-CI), and manually counted with a hemocytometer (Bright-Line Z359629). Mono- and co-cultures of each subclone were seeded across a range of initial relative proportions in 96-well formats and allowed to attach for 18-24 hours.

Wells were treated with the following drugs: gefitinib, paclitaxel (Cayman 10461), etoposide (Cayman 12092), pemetrexed (Cayman 26677), and lapatinib (Cayman 11493) as single agents. Plates were loaded into a BioSpa 8 Automated Incubator (BioTek Instruments). Time-lapse microscopy images were obtained for bright field, GFP, and mCherry via Cytation 5 Imaging Reader (BioTek) every 4 hours over the course of 5 days.

### Image Processing

Images were processed with Gen5 (BioTek) and the open-source software ImageJ.^49^ Image sets were duplicated, background subtracted, contrasted limited adaptive histogram equalization (CLAHE), and thresholded. Despeckle filter was applied to the now binary images, watershed segmentation was performed, and raw cell numbers were extracted from the resulting image sets.

### Evolutionary Game Assay

To quantify the dynamics in our *in vitro* environments, we utilized the experimental game assay developed by Kaznatcheev et al..^30^ Initial proportions were calculated for each well individually from the first image. Time series of raw cell numbers were normalized against initial number in each well. Linear regression was performed using the Theil-sen estimator on the semi-log cell change against time. The slope of the resulting linear function (with its corresponding 95% confidence interval) was translated as the growth rate across the time series, which were normalized against the average of six parental monoculture wells that were run on each plate.

To find the dependence of fitness on the frequency of subclonal interaction, least squares regressions were performed on the growth rate against the initial proportion of parental in each well. This regression was weighted against the inverse of the errors 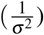 associated with each growth rate. The resulting linear equations describe fitness as a function of the initial proportion of the opposing subclone:

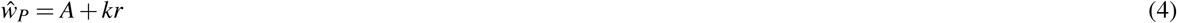

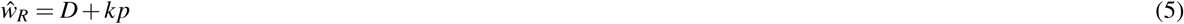

The intercepts of these functions translate to monoculture fitness, which are the symmetric payoffs within a game matrix. The asymmetric payoffs can be translated as the fitness values when *r* and *p* are equal to 1:

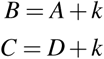

These linear equations can be rearranged to describe the fitness (*ŵ*) of a sub clone as a function of the initial proportion (*p*) of interacting cells within the population.

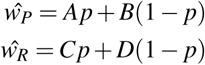

Payoff matrices corresponding to each of the different conditions can be derived by setting *p* equal to one and zero for both equations. For example, the symmetric payoff for parental occurs when *p* = 1, which translates to *ŵ*_*P*_ = *A*.

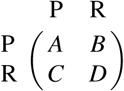

The errors associated with the on-diagonal payoffs are equivalent to the uncertainty of the intercept values, *σ*_*A*_ and *σ*_*D*_ for parental and resistant respectively. The errors associated with the off-diagonal payoffs were derived by propagating the uncertainty of both the intercept and slope through both the intercept and slope of (**eq. 6**):

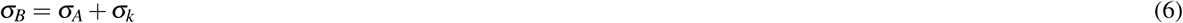

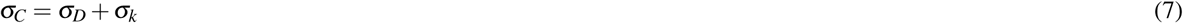

### Growth models

To synthesize hypothetical tumor growth using our measured frequency-dependent growth rates, we used two distinct models, one that allowed for infinite growth and one that limited total volume to a strict maximum. This was done to identify salient qualitative features that persisted across this spectrum of models, rather than make specific quantitative predictions.

For infinite growth, replicator dynamics were chosen:

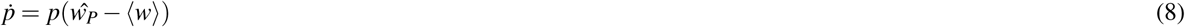

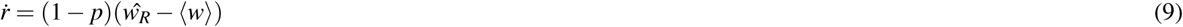

where ⟨*w*⟩ denotes average population fitness (i.e., *w* = *pŵ*_*P*_ + (1 *− p*) *ŵ*_*R*_).

For growth that is strictly limited to a maximum, a Lotka-Volterra derivative^50^ was utilized that included frequency - dependent growth:

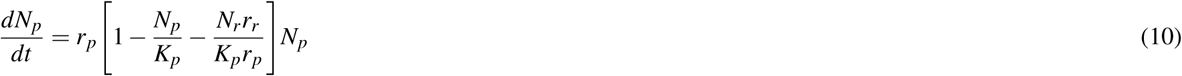

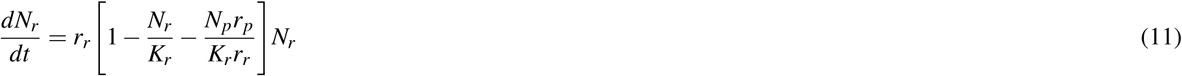

where *r*_*p*_ and *r*_*r*_ are non-cell autonomous growth rates determined by values of the game matrix such that:

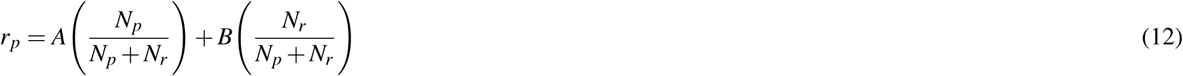

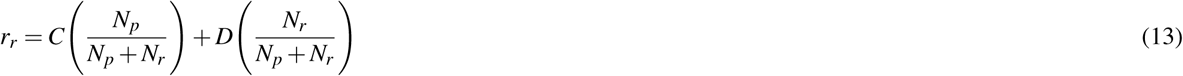

While this model is insensitive specific carrying capacity values, it is highly sensitive to the relative value of the carrying capacity. Given that both subclones occupy similar space in an *in vitro* environment, we first evaluated the condition where the carrying capacities were equal to one another (*K*_*p*_ = *K*_*r*_). This assumption likely does not translate to *in vivo* conditions.

Instead, the carrying capacities of each type would likely vary across different environments. To capture this phenomenon, the carrying capacity was also scaled for each condition:

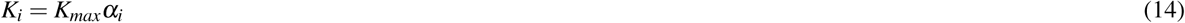

where *K*_*max*_ is the maximum carrying capacity across all conditions and *α*_*i*_ is a weighting term that scales this maximum using a ratio of monoculture growth rate in the current condition against the maximum growth rate 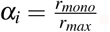.

